# Out-of-sight partners: autonomous wearable supernumerary limbs for dynamic co-manipulation

**DOI:** 10.64898/2026.09.02.748789

**Authors:** Dorian Verdel, Bjorn Xu, Hector Cervantes Culebro, Alexis Devillard, Luca Mattioli, Dario Farina, Philippe Souères, Etienne Burdet

## Abstract

Supernumerary robotic limbs (SLs) could extend human motor capabilities beyond the natural body, yet most demonstrations have been limited to simple, quasi-static tasks or direct teleoperation under visual control. Whether humans can safely and intuitively collaborate with autonomous SLs during complex, dynamic tasks remains unknown. We studied this question in a demanding assembly task requiring coordinated transport and handovers of a long object, which could not be completed by a single person. We developed a reconfigurable backpack platform mounting up to four high-payload robotic arms, together with safety-aware motion planning that generates human-compatible trajectories from task states without training data. Despite the SLs operating out of sight, participants adapted quickly, completed the task reliably, and reported low cognitive demand alongside high perceived safety and predictability. Kinematic and force analyses showed that control design governed both movement fluency and whole-body stability: faster limb motions yielded smoother interactions and shorter completion times, and a mirrored coordination strategy–in which one robotic limb counterbalanced the other–further shortened duration and reduced variability in left-right ground reaction force differences during co-manipulation. These results show that humans can integrate autonomous wearable limbs as out-of-sight partners in dynamic whole-body collaboration, and delineate behavioral and control principles underpinning fluency, stability, and usability in human augmentation.

## 1 Introduction

Movement augmentation seeks to extend human action beyond the abilities of the natural body. Among the most direct approaches are *supernumerary robotic limbs* (SLs), wearable effectors that add new manipulation capabilities by complementing *natural limbs* (NLs). Such devices could transform how humans interact with the physical world by enabling a single operator to perform tasks that would otherwise require additional people, expanding the reachable workspace, and supporting users with motor impairments [1, 2]. Recent progress has shown that supernumerary limbs can assist grasping, stabilization, and simple manipulation [3, 4]. However, most demonstrations remain confined to quasi-static scenarios, direct teleoperation, and task-specific systems in which the robotic limbs are visually monitored and only weakly coupled to whole-body movement.

A critical open question for real-world augmentation is whether humans can safely and intuitively coordinate with autonomous wearable SLs during complex dynamic tasks. In practical settings, wearable robotic limbs will often operate outside the user’s visual field, share workspace with the NLs, and interact through manipulated objects that mechanically couple SLs and NLs. Under these conditions, successful augmentation depends not only on robotic control accuracy, but also on how humans perceive, predict, and adapt to the behavior of the added limbs. Autonomous control must therefore satisfy several requirements simultaneously: it must remain safe and legible [5], adapt to the user’s posture and ongoing actions to support whole-body balance [6, 7], and reduce rather than increase the cognitive burden associated with supervising extra degrees-of-freedom.

Progress toward these objectives has been limited by both hardware and control constraints. On the hardware side, many supernumerary limb systems have been designed for narrow applications, such as part stabilization for assembly [8–10], augmented manipulation with additional fingers [11–15], balance improvement [16, 17], or artistic performance [18, 19], making it difficult to study dynamic whole-body coordination across tasks. On the control side, prior work has often relied on direct control via body interfaces [15,20–27], learning from demonstration [8,28], or task-specific heuristics. Although effective in constrained conditions, these approaches do not fully address the demands of dynamic co-manipulation, in which the robotic limbs must plan movements that are predictable to the user–sometimes outside visual supervision–and often under biomechanical and balance constraints. As a result, it is still unclear whether autonomous wearable limbs can function as true collaborative partners rather than as tools that require continuous user oversight.

Here we address this gap by studying human coordination with autonomous out-of-sight wearable limbs during a dynamic whole-body task. We developed an integrated experimental framework combining a reconfigurable backpack platform [29], carrying two to four high-payload robotic arms, with a safety-aware motion-planning architecture that generates human-compatible trajectories from task states without training data. We tested this framework in a demanding assembly task requiring coordinated transport and handovers of a long object across the body, a task that cannot be completed by a single person and in which SLs operate outside the user’s field of view during critical phases.

Using kinematic, force-plate, and perceptual measurements in participants augmented with two back-mounted SLs, we asked three questions. First, can humans rapidly adapt to autonomous SLs and collaborate with them reliably when they cannot be visually monitored? Second, how does control design affect the fluency of co-manipulation? Third, can the coordination of several SLs improve whole-body stability during dynamic interaction? Our results show that participants rapidly integrated the autonomous limbs into task execution, reported low cognitive demand and high perceived safety and predictability, and completed the task reliably despite the limbs being out of sight. We further show that faster SLs motions improved fluency and reduced task duration, while a mirrored coordination strategy, in which one robotic limb counterbalanced the other, improved balance-related outcomes during co-manipulation. Together, these results establish a foundation for understanding how autonomous wearable limbs can be integrated into human motor behavior and identify control principles for safe, fluent, and usable human augmentation.

## 2 Results

### 2.1 An environment to investigate human movement augmentation

The *MUlti-limb Virtual Environment* (MUVE) for full body and multisensory (visual, audio, haptic) interactions is a unique system enabling physical interaction with arms, legs, and supernumerary robotic limbs in dynamic virtual environments (Supplementary Fig. S.1A, [29]). MUVE integrates the Motek GRAIL system [30], comprising (i) 3D motion capture (Vicon), (ii) immersive 3D virtual reality, and (iii) a split treadmill-based VR motion suite integrating force plates, as well as (iv) up to four lightweight, wearable robotic arms for movement augmentation. Critically, all components are time-synchronized and integrated within a single system using ROS.

We require generic SLs that are (i) safe to use, (ii) versatile across tasks, (iii) simple and fast to reconfigure, (iv) that provide a workspace commensurate with the operator’s natural arms, (v) offer a high payload, and (vi) support multiple control modalities. Because dedicated SLs developed to date [3, 4] do not satisfy all of these requirements and are not widely available, we turned to commercial collaborative robots (cobots). Most current cobots (e.g., KUKA LWR, Franka Emika Panda, UR7e Universal Robot) offer payloads of 3–7.5 kg but weigh 16–20.6 kg and their reach of 79–85 cm may not be sufficient for coordinated operation with the human arms. The Unitree Z1 weighs only 4.5 kg with a 3 kg payload, but its base joint actuator cannot sustain the full arm moment when extended horizontally, and its reach of 74 cm with 6 axes is too limiting to collaborate well with the user’s arms.

By contrast, the Kinova Gen3 has link lengths exceeding those of human arms and a 89.1 cm reach (Supplementary Fig. S.1B), offering a sufficiently large workspace when back-mounted to carry out co-manipulation with the operators’ NLs (Fig. 1A,D). Its 7 *degrees-of-freedom* (DoF) afford suitable manipulability (Fig. 1B), and its mass of 8 kg with a 4 kg payload is attractive. We therefore selected four Kinova Gen3 arms as SLs within the MUVE.

**Figure 1:**
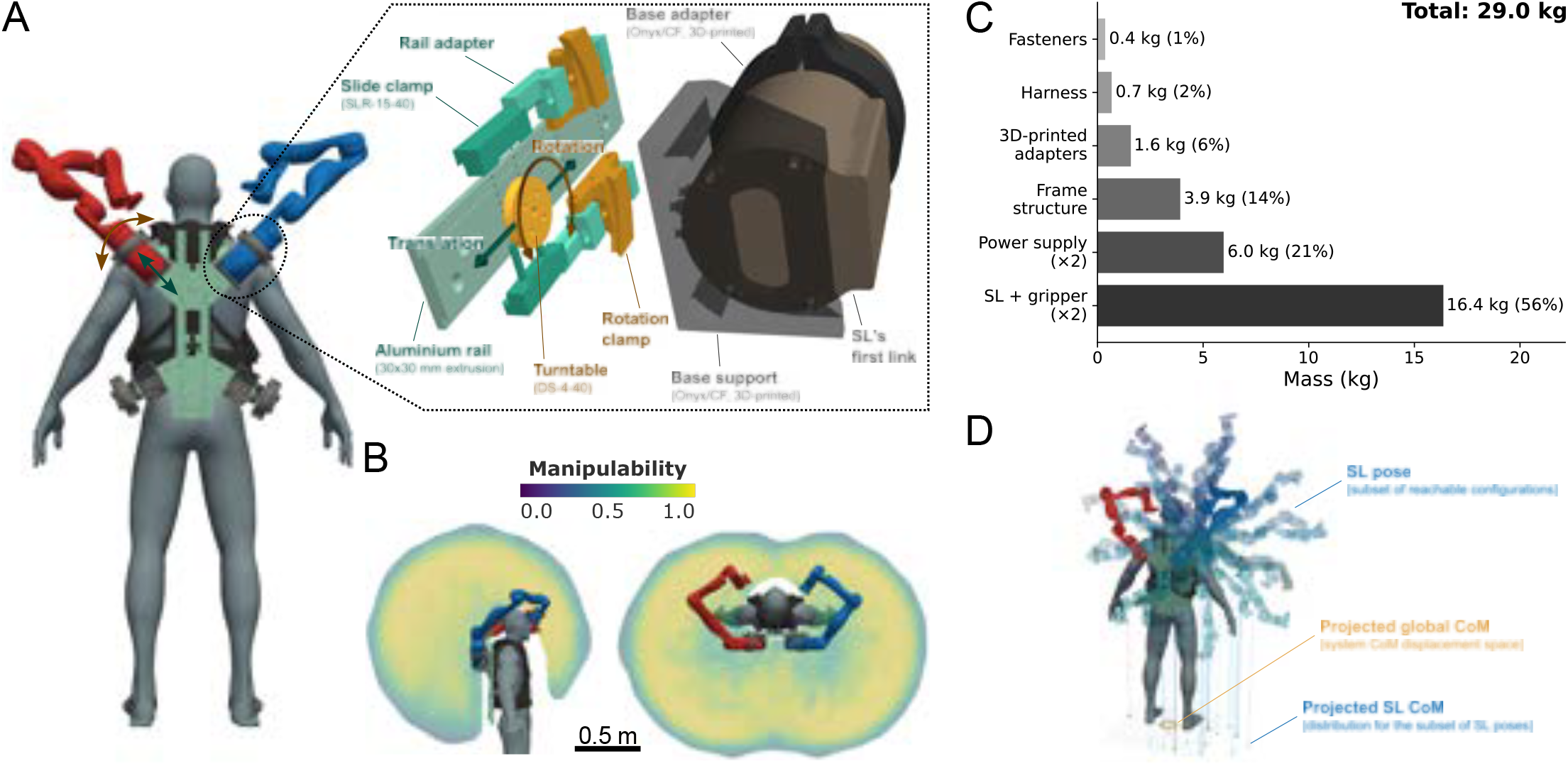
Design of the modular backpack for SLs. **A.** The backpack allows the operator to carry up to four SLs: two with bases facing upward and two facing downward. The orientation and position of each SL base on the frame can be adjusted to meet task-specific requirements. **B**. “Dr. Hexapus” configuration with two upwards-facing SLs (here two Kinova Gen3 arms as in our validation experiment), providing a large shared workspace: it overlaps the user’s natural reach for collaboration with the NLs, and extends to regions behind the user that are otherwise unreachable, enabling object retrieval and co-manipulation of large items. In all those regions, it provides a high manipulability as assessed using Yoshikawa’s index [31]. **C**. Mass of the backpack components. **D**. SL movements within the workspace can substantially shift the combined human–SL center of mass (CoM), motivating coordination strategies that promote stability.

To enable augmented manipulation, we developed a “Dr. Octopus” backpack: a reconfigurable research platform that enables rapid changes in SL configuration by varying the number of SLs (2 or 4) and their positions and orientations, without redesigning the support or performing complex operations. The backpack consists of a rigid X-shaped frame that secures the SLs on the user’s back, coupled with an industrial vest (Stihl RTS, Waiblingen, Germany) that distributes the load across the shoulders and hips.

Modularity is achieved at the interface between each SL base and the rigid frame. Each SL is clamped to a rail via a Montech DS-4-40 turntable and SLR-15-40 sliding element, providing two continuous adjustment axes (see Fig. 1A): translation along the rail and rotation about the rail normal. Reconfiguring the setup—for example, moving an SL from the upper rails to the lower rails—requires only loosening the rail clamp with a simple key and takes under five minutes.

For the complex task presented in Section 2.3, we mounted two SLs on the upper rails, approximately symmetric about the user’s spine. As shown by the manipulability analysis in Fig. 1B, this arrangement supports a large shared workspace that enables versatile human–SL and SL–SL physical collaboration. As reported in Fig. 1C,D, the backpack mass in this configuration and the effect of SL postures on the combined human–SL *center of mass* (CoM) are important considerations for controller design, motivating the inter-SL coordination strategies detailed in Section 4.4.

### 2.2 Human-SLs interaction task

To evaluate human-SLs interactions, we asked 8 participants to perform a complex collaborative assembly task while using two SLs that autonomously compute safe action goals and movements based on the operator’s real-time state and a predefined sequence of actions. The task was designed to: (i) require more than two arms (thus it cannot be completed by a single operator), (ii) involve several types of NLs-SLs coupling, (iii) impose accuracy constraints, and (iv) preclude visual monitoring of the SLs. The selected task comprised seven actions to move a 3 m long PVC tube of mass 1.015 kg from the participant’s right side to a target assembly zone on the left, where it was screwed to a support (Fig. 2A–C). Importantly, holding the tube with one hand would induce a torque of roughly 14.7 Nm at the wrist, preventing its lone manipulation and necessitating co-manipulation with an SL.

**Figure 2:**
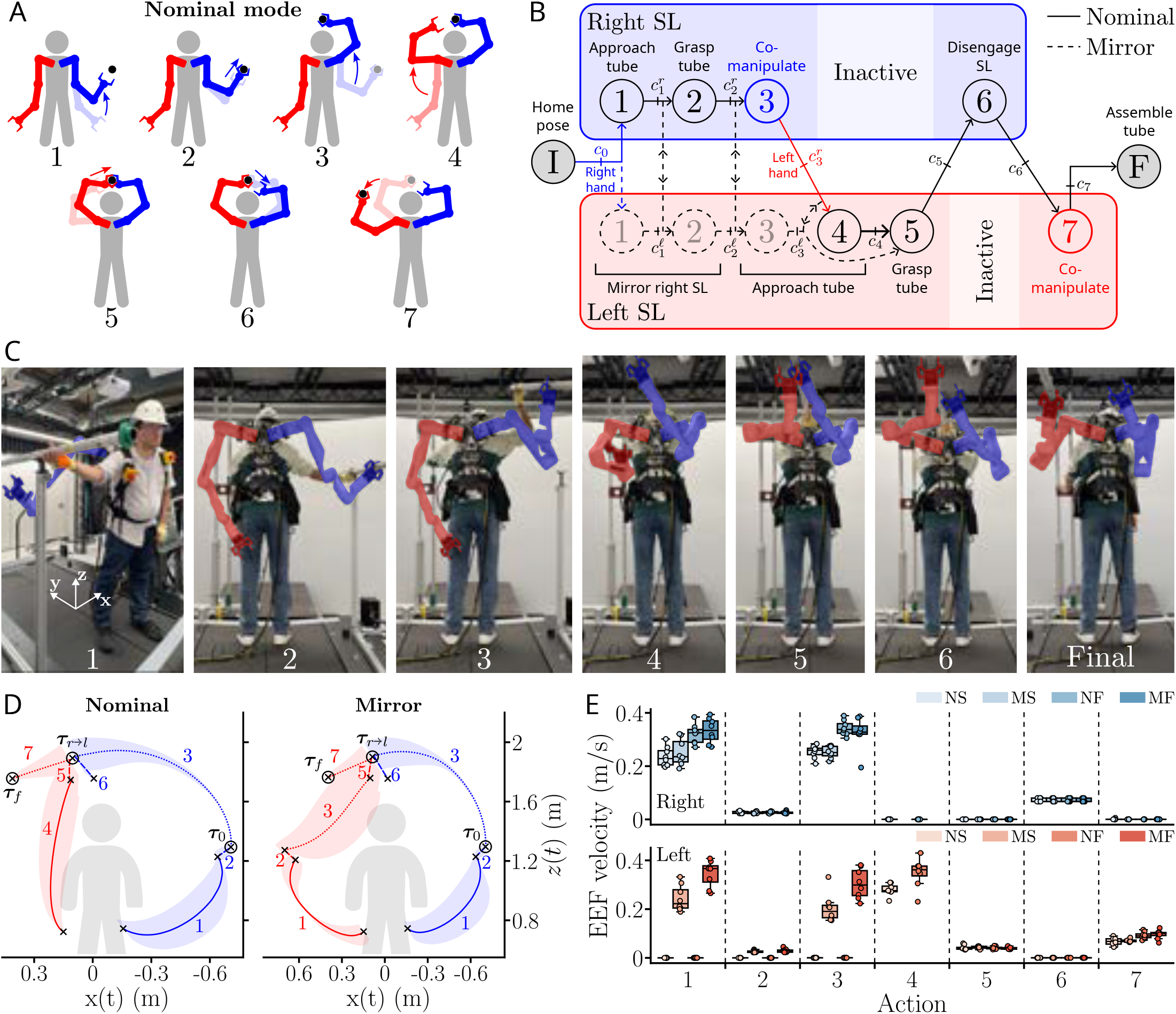
Setup and assembly task for augmented worker. Photographs are of the authors. **A**. The task was decomposed into seven actions to grab, transfer and assemble the tube. In the *nominal mode*, actions were executed sequentially; in the *mirror mode*, the left SL mirrored the right SL movements during actions 1 and 2, and performed action 3 in parallel to the right SL. **B**. The assembly task was modeled as a finite state machine with conditional transitions. Goals were joint configurations realizing a desired 6D EEF pose, computed via inverse kinematics. In *nominal mode* (solid lines), the controller reached the right SL intermediate goals **g**^*r*^ (poses in task space with *i* action index) to satisfy conditions 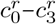. In *mirror mode* (dashed line), both right and left SLs had to reach their intermediate goals before transitioning to action 5. Supplementary videos illustrate the task execution in both modes. **C**. Participants wore two back-mounted SLs to perform the collaborative assembly, here shown for the nominal mode. **D**. Average trajectories of the SLs EEFs in nominal and mirror modes, with right SL data in blue and left SL data in red. Tube poses ***τ***_0_ (initial), ***τ***_*r*→*l*_ (right-to-left handover), and ***τ***_*f*_ (final assembly pose). Crosses represent goals of SLs movements that, when reached, allowed to validate state transition constraints. **E**. Average SL’s EEF speed in each action for the four tested conditions: {nominal–slow (NS), mirror–slow (MS), nominal–fast (NF), mirror–fast (MF)}.

Transferring the tube from right to left required coordinated handovers of the operator and the SLs above the head, followed by the lateral displacement. In details, the task was decomposed into the following actions (Fig. 2A–D): (i) prepare to grasp tube from support with right SL, (ii) grasp tube with right SL, (iii) *co-manipulate* the tube with right effectors (SL and NL), (iv) move left SL to prepare handover over human head, (v) grasp tube with left SL, (vi) disengage right SL from tube, and (vii) *co-manipulate* the tube with left effectors (SL and NL). Actions involving grasping or positioning of tube for assembly imposed accuracy constraints requiring to account for each participant’s anthropometrics, posture and behavior, rendering the task planning complex and unsuitable for offline computation. The *tube approach actions* of the SLs (actions 1& 4) were initiated when the participant’s right- and left-hand got close enough to the tube, respectively. Planning and execution of actions 2& 3, and 5–7 were initiated upon completion of the preceding action. This action sequence and its triggers were formalized as the state-transition graph in Fig. 2B for the nominal and mirror control modes detailed below.

Maintaining the operator’s balance is critical when working with worn SLs, whose motions can produce significant shifts in the combined human-SL CoM [6, 7]. In addition to the (i) *nominal control mode*, in which the two SLs executed the seven actions in sequence (Fig. 2A–C), we tested a (ii) *mirror control mode*, in which the left SL movements were symmetric to the right SL’s movements during the first two actions and performing action 3 in parallel to the right SL, to counteract the right SL’s mass and reduce lateral destabilization. We also tested two velocity levels: (i) *fast*, near the maximum feasible speed according to the robots capabilities, and (ii) *slow*, approximately 30% slower. Those two velocity levels allowed us to (i) test whether increased reaction torques applied on the human in the *fast* condition hampered safety, and (ii) analyze how SLs’ movement speed impacts the operator behavior.

These factors yielded four conditions: {*nominal-slow* (NS), *nominal-fast* (NF), *mirror-slow* (MS), *mirror-fast* (MF)}. Each participant performed three collaborative assembly trials per condition. To assess human-SL interactions throughout task execution, we synchronously recorded: (i) kinematics using SLs embedded encoders and an optoelectronic motion capture system, and (ii) *ground reaction force* (GRF) at the feet using one force plate per foot. We complemented these measurements with questionnaires to obtain a comprehensive evaluation of performance and perception while wearing SLs.

### 2.3 Human-SLs performance

Fig. 2D illustrates the SLs behavior in the nominal and mirror modes. SL trajectories are smooth and broadly repeatable (Fig. 2D, Fig. 3A,B), with variations arising from anthropometric differences across the participants, and from differences in participant positioning within the scene, both of which are accommodated online by our motion-planning algorithm. As expected, in mirror mode, actions 1–3 are also performed by the left SL, unlike in nominal mode where the SL movement goes straighter towards the *τ*_*r*→*l*_ goal after the end of the right SL movement. Fig. 2E shows how the commanded movement velocities affected average EEFs speeds (see statistical analyses in the Supplementary Materials), demonstrating that users can successfully complete our complex task across a wide range of commanded speeds while wearing the SLs and co-manipulating the tube. Higher speeds and co-manipulation actions tended to produce larger SL joint torques (Supplementary Fig. S.2), suggesting that the SLs’ power-to-weight ratio may quickly become a limiting factor for sustained physical interactions.

**Figure 3:**
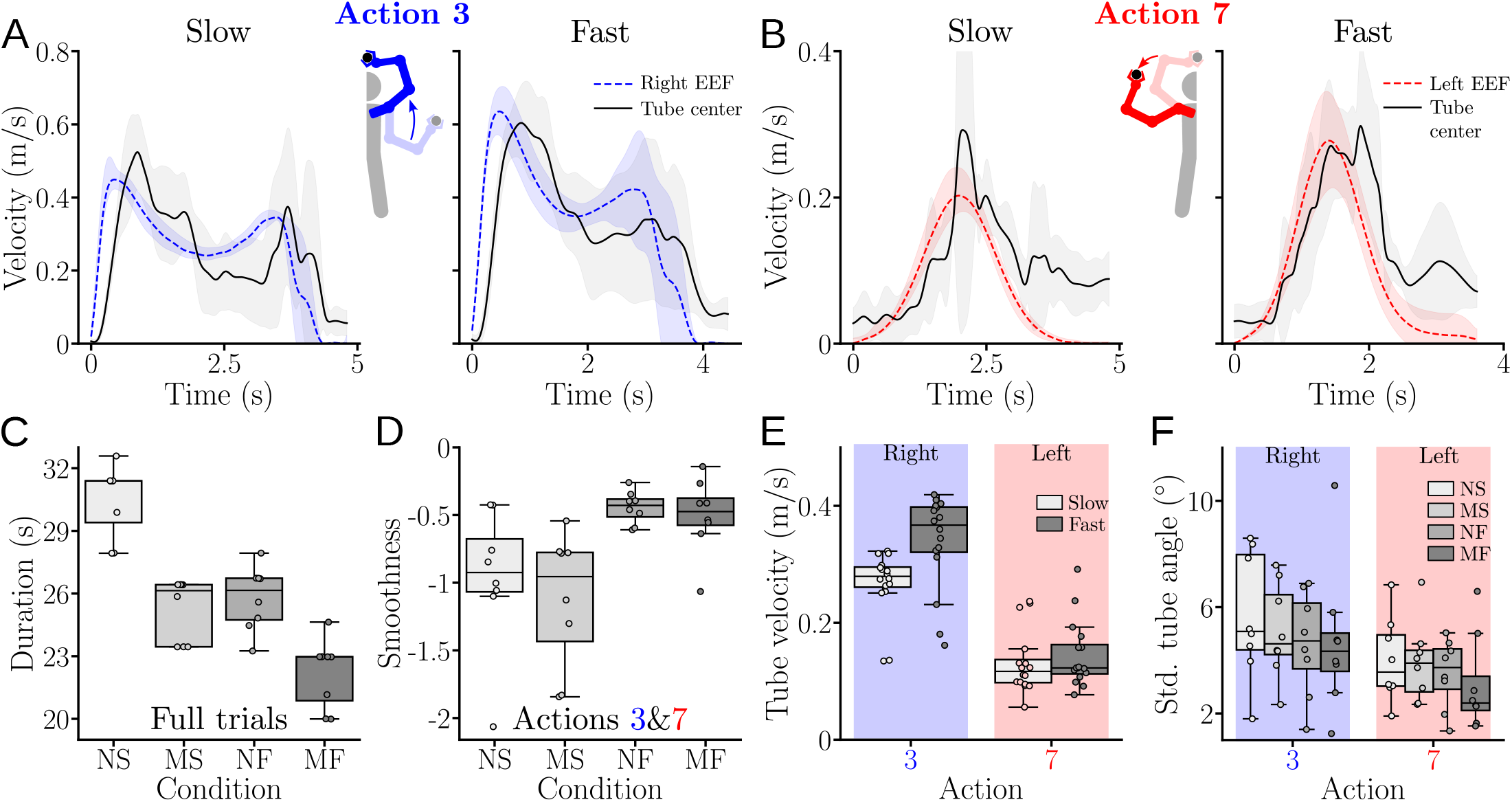
Kinematics of NL-SL co-manipulation. Data related to the right SL are in blue, to the left SL in red, and to the tube in black. **A**,**B**. Mean velocity profiles of the right and left EEFs and the tube during action 3 and 7, respectively. **C**. Total duration of the seven actions across control modes. **D**. Tube movement smoothness during co-manipulation across conditions. **E**. Mean tube speed under fast vs. slow SL control; nominal and mirror modes data were pooled by velocity for clarity. **F**. Standard deviation of the tube orientation in actions 3 and 7 for all four conditions.

To further understand human adaptation to the SLs, we analyzed user behavior across control conditions. First, we examined task and tube kinematics during co-manipulation, summarized in Fig. 3. The average velocity profiles of the right and left *endeffectors* (EEFs) exhibit smooth motion during *transport phases with co-manipulation* (actions 3 and 7) in slow and fast conditions, with shorter duration and higher peak velocities in fast trials (Fig. 3A,B). The tube velocity profiles follow a similar shape to the SLs’ EFF profiles, with higher velocities in the fast conditions, and a lag. Overall, participants appeared to adapt their movement patterns to those of the SLs. Notably, during action 3 where the right SL optimal control yielded a velocity profile with two peaks, the human adopted a similar strategy, different from the bell-shape velocity profile generally adopted in point-to-point movements.

Quantitatively, we observed reduced movement time in mirror mode (Fig. 3C) (*W* = 0.93, *p <* 0.001, main effect of control mode on the task duration). Fast conditions {NF,MF} led to shorter durations than their slow counterparts {NS,MS} (*p <* 0.008, *D >* 1.92 in both cases). Furthermore, both slow and fast mirror conditions {MS,MF} led to shorter durations than their nominal counterparts {NS,NF} (*p <* 0.01, *D >* 2.24 in both cases). This shows that the mirror condition reduced task duration, in addition to potentially increasing stability.

We then conducted a smoothness analysis using dimensionless jerk [32] (Fig. 3D), which revealed a main effect of condition when grouping the transport actions 3 and 7 (*W* = 0.64, *p* = 0.001). Pairwise comparisons showed no effect of control mode (nominal v.s. mirror), with mirror conditions exhibiting slightly greater variability. However, the fast conditions lead to smoother movements with less jerk variability than slow conditions (*p <* 0.025, *D >* 1.1 in all cases). In sum, faster SL movements yielded smoother interactions, with no clear impact of control mode.

To further characterize human–SL coordination, we analyzed tube motion during co-manipulation (Fig. 3E,F). Because there was no significant difference between the velocity in the nominal and mirror modes, we grouped trials in these two modes, and observed that the average tube velocity in actions 3 and 7 was higher in fast vs. slow conditions (Fig. 3E; *W* = 1, *p <* 0.005, main effect of velocity). Pairwise comparisons showed this difference to be significant for action 3 only (*p <* 0.006, *D* = 1.06), with action 3 yielding significantly faster movements than action 7 overall (*p* = 0.016, *D* = 2.77). Beyond the mean speed, we assessed human–SL coordination through the variability of tube orientation in the horizontal plane (Fig. 3F). This variability was lower in action 7 than 3 (*p <* 0.01, *D* = 3.58), with no clear effect of the condition. Finally, a Granger-causality analysis indicated that SL EEF movements preceded and predicted tube motion in both action 3 (*F >* 19.4, *p <* 0.001 in all four conditions) and action 7 (*F >* 98.65, *p <* 0.001 in all four conditions). Overall, this indicates that participants adapted their movement velocity—both in magnitude and profile—to SL behavior, and human control of the tube tended to lag SL movements.

To analyze the effects of human-SLs interaction on human balance, we analyzed the ground reaction forces induced by the right and left foot of the user with the different control modes. (Fig. 4). The GRF ellipses for the first three actions in the (*x, z*)–plane, reported in Fig. 4A, suggest that there is no large difference in GRF intensity between the nominal and mirror modes, but the GRF in the nominal mode tended to be more inclined than in the mirror mode, especially in actions 1 and 2. Overall, the GRF ellipses, and further analysis of the difference of GRF between the two feet described below, suggest that the mirror mode led to a more balanced human-SLs system during action 1, showing the importance of counter-weighting SLs’ displacements during free movements.

**Figure 4:**
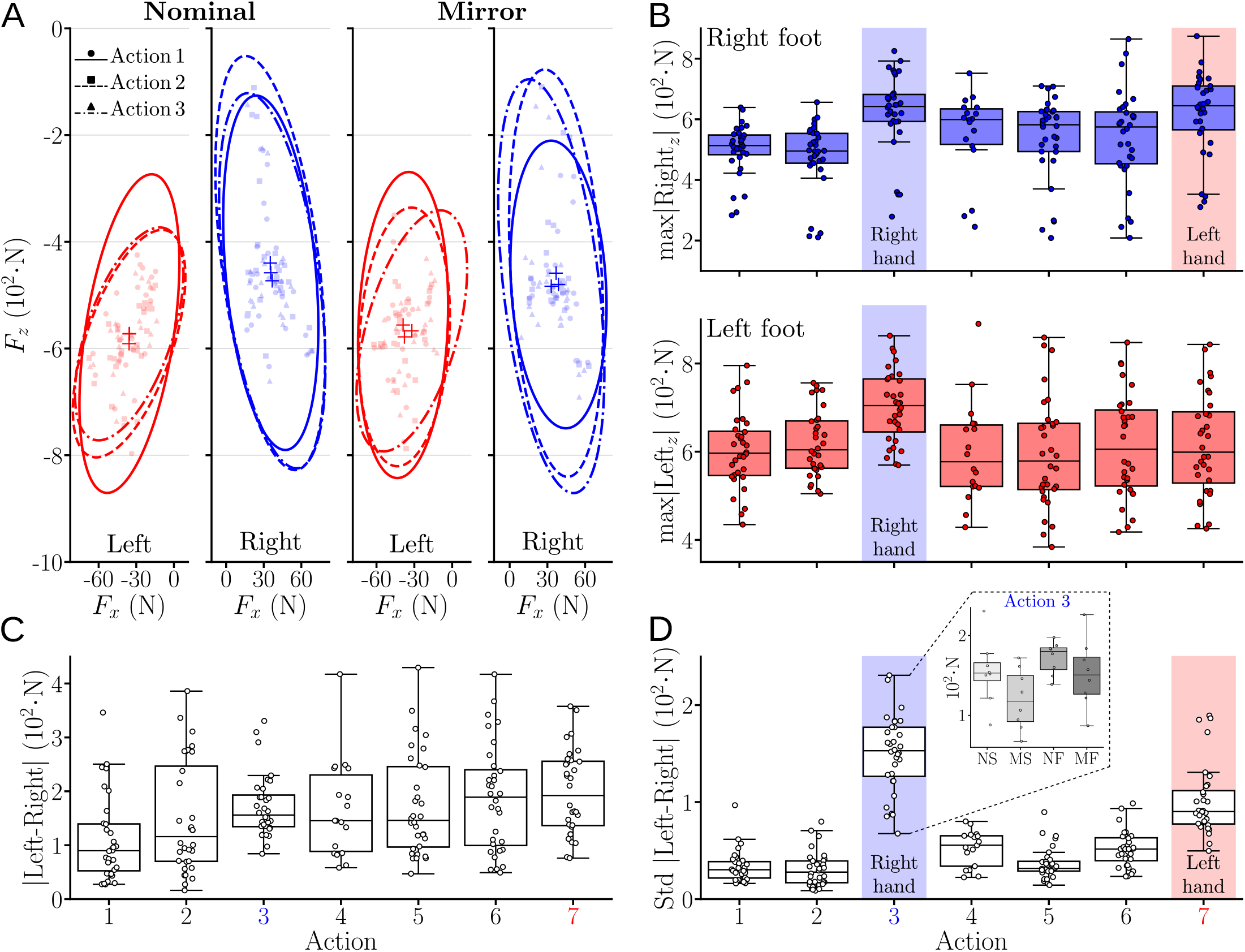
Ground reaction forces during human-SLs collaborative assembly. Right-side hand and foot data are shown in blue, left-side data in red. Results are reported for each action; nominal and mirror modes are pooled across conditions for readability, with detailed analyses in the text. Co-manipulation actions are highlighted with labeled shaded bands when they exhibited significantly higher effort or variability. **A**. Distribution of left and right GRFs in actions 1–3 for the nominal and mirror modes. **B**. Absolute maximum of the vertical GRF component for the right (top) and left (bottom) foot across the seven actions. **C**. Absolute difference between the left and right GRF. **D**. Standard deviation of the absolute left-right GRF difference; inset shows between-condition differences in co-manipulation action 3.

Furthermore, the performed action affected the vertical GRF magnitude with a main effect for both the right foot (*W*_6_ = 0.65, *p <* 0.001) and left foot (*W*_6_ = 0.43, *p* = 0.002), as illustrated in Fig. 4B. Specifically, actions 3 and 7 yielded higher vertical forces on the right foot than all other actions (*p <* 0.049, *D >* 0.5 in all cases), and action 3 yielded higher vertical forces on the left foot than all other actions (*p <* 0.033, *D >* 0.92). In sum, dynamic co-manipulation of the tube by the human and SLs increased vertical loading.

To examine potential lateral balance biases (i.e., horizontal displacements of the human-SLs CoM), we analyzed the averaged difference of GRF between the left and right feet. Fig. 4C shows that this difference was mainly action-dependent (*W*_6_ = 0.28, *p* = 0.039, main effect of the action). Furthermore, the mirror mode tended to produce slightly lower balance biases than the nominal mode primarily in actions 1 & 3. This reduced bias was confirmed by a main effect of the control mode in action 1 (*W*_3_ = 0.37, *p* = 0.03), with lower bias in MF than NF (*p* = 0.047, *D* = 0.7). This cannot be observed in actions 5–7 that were not changed by the control mode and there is no action 4 in mirror mode. In summary, the mirror mode tended to reduce the lateral balance bias relative to the nominal mode, although this effect was modest and statistically confirmed only in action 1.

Finally, we analyzed the variability of the left–right vertical GRF difference within each action to assess whether the combined CoM oscillated about its mean position, which would indicate greater instability. Fig. 4D shows that dynamic co-manipulation of the tube in actions 3 & 7 increased this variability, which was confirmed by a main effect of the action on the difference variability (*W*_6_ = 0.76, *p <* 0.001). In details, the variability was higher in action 3 than in all the other actions (*p <* 0.023, *D >* 2.29 in all cases), and higher in action 7 than in all actions except action 3 (*p <* 0.015, *D >* 2.13 in all cases). We also observed a main effect of control mode on the variability of the GRF difference for actions 1 (*W*_3_ = 0.51, *p* = 0.006) and 3 (*W*_3_ = 0.37, *p* = 0.03). In action 1, condition MF yielded better stability than NF (*p* = 0.047, *D* = 0.89), and a non-significant but large improvement relative to NS (*p* = 0.07, *D* = 0.79). In action 3, MS yielded better stability than NF (*p* = 0.047, *D* = 1.71) and overall variability was substantially lower in mirror than nominal mode (*p* = 0.05, *D* = 1.27). Overall, this shows that the dynamic co-manipulation of the tube increased the instability of the human-SLs system, which was attenuated by mirroring the movements of the SL carrying out the operation.

### 2.4 Ergonomics of human-SLs interactions

To complement the objective assessments above, we evaluated overall ergonomics using a questionnaire inspired by NASA-TLX [33]. The three results on difficulty in Fig. 5 first indicate that assembling the tube was physically demanding for most participants, but not especially mentally taxing or difficult to understand. Next, three comfort-related items showed that the backpack felt uncomfortable and exerted pressure on the shoulders, likely due to its relatively high mass. Finally, two items on predictability and safety revealed that, despite being unable to see the robots during the task, most participants could anticipate SL behavior and felt safe during the collaborative assembly. Overall, participants identified physical effort and comfort as the primary drawbacks, whereas SL use was not experienced as cognitively demanding or perceived as unsafe.

**Figure 5:**
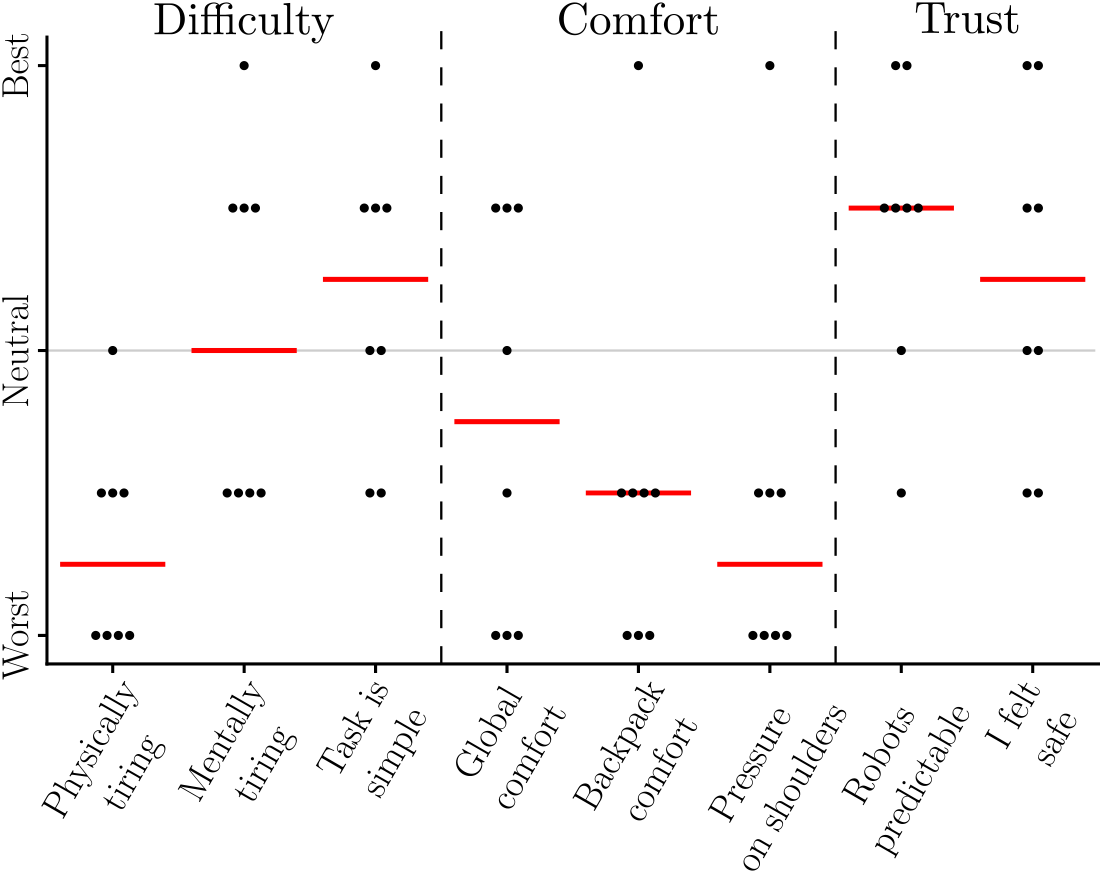
Participants’ perception of the task and SL interaction. Each point corresponds to one participant; red horizontal lines indicate the median across participants.

## 3 Discussion

We showed that people can rapidly collaborate with autonomous, out-of-sight supernumerary robotic limbs (SLs) to execute a demanding whole-body task that a single person cannot perform alone. Participants reliably coordinated transport, handovers, and assembly of a long object with the SLs without seeing them, reporting low cognitive demand, high predictability and safety. Across the tested conditions, faster SL motions produced smoother interactions and shorter completion times, and a mirrored coordination strategy–using one SL to counterbalance the other during co-manipulation–reduced task duration and improved balance-related outcomes. These results yield concrete hardware and control principles for fluency, stability, and usability in dynamic augmentation.

Prior robotic augmentation studies have mostly targeted quasi-static, visually supervised scenarios with weak coupling to natural limbs (NLs), including tasks with SLs focused on static holding or drilling with negligible balance effects [8–10, 28, 34], and supernumerary fingers offering performance far from polydactyly capabilities [15, 35, 36]. By contrast, we provide one of the first demonstrations–and quantitative analyses–of complex, whole-body, dynamic co-manipulation with autonomous SLs operating operating outside the user’s visual field, measuring interaction fluency, human adaptation, and balance.

To enable these advances, we developed a one-of-a-kind modular platform that enabled in-depth investigation of human behavior when interacting with SLs. The reconfigurable backpack we designed can mount up to four SLs and quickly adjust their base pose to suit different tasks. The system provides high manipulability within a large shared workspace, both in front of the user (enabling close collaboration with the NLs) and behind the user (extending reach to otherwise inaccessible regions). Its main limitation is mass (≈ 30 kg with two SLs), perceived as physically demanding at shoulders and hips. Here, we prioritized experimental versatility and reproducibility by using a modular platform built around commercial cobots with high payload. The same platform can also host custom, lighter arms (e.g., carbon-fiber links) or offload backpack weight via supernumerary legs [37, 38].

To minimize cognitive demands, we implemented autonomous behaviors within a versatile human–SL interaction framework [2] that: (i) ensures safety via collision avoidance within and between SLs and with the user and environment; and (ii) coordinates limbs via a mirror mode that uses the second SL as a counterweight. Motion planning with integrated collision avoidance was performed via trajectory optimization [39] rather than deep-learning or probabilistic methods [40, 41], yielding legible motions without training data, and providing clearer planning guarantees on safety and biomechanical compatibility, both of which are critical for human-SLs interactions.

Safety can be implemented at several levels. Hard workspace bounds encoded as inequality constraints offer strong guarantees [42–44], but are difficult to implement and computationally expensive with nonlinear, time-varying constraints. Dynamics-level repulsive force fields are simpler to implement and computationally efficient [45], yet can yield unpredictable, inefficient paths when multiple fields from human NLs and the environment interact during movement. We therefore adopted an intermediate approach, by penalizing the signed distances to obstacles in the cost function using voxelized environment models [46]. This approach can be adapted depending on the application, e.g. with fine voxelization in tight workspaces and coarse voxelization in open scenes, and could be extended with multiscale voxelization for time-varying occupancy maps to penalize predicted conflicts.

Participants successfully collaborated with their SLs to complete the complex assembly task, reporting low cognitive load and high perceived safety and predictability. They rapidly adapted NL motion to SL behavior, modulating the shape and magnitude of the tube’s velocity profile according to the action and the SL control mode. Crucially, this occurred without visual monitoring or additional artificial sensory feedback (e.g., vibrotactile cues) [47], indicating that the co-manipulated object provided a robust haptic communication channel [48–50]. Mechanical coupling through the tube directly conveyed interaction forces, enabling users to anticipate and regulate SL behavior from low-dimensional hand cues while avoiding the neural resource-allocation and sensory feedback learning burdens associated with explicit sensory substitution or continuous visual monitoring [51, 52].

The adaptation patterns suggest that participants did not treat the SLs as external perturbations, but as partner-like agents in a shared control loop. During co-manipulation, human movements aligned with SL behavior, and a Granger-causality analysis showed that SL motions preceded and predicted object motion. This means that users did not supervise every robotic movement, but instead they leveraged the robot’s predictability to collaborate. Effective augmentation may therefore depend less on maximizing the robot responsiveness than on establishing stable, intelligible interaction dynamics that users can rapidly internalize and incorporate into their own motor control. Consistently, SL motions slower than typical human timing degraded movement smoothness and increased task duration, in line with theories that faster timing reflects minimization of an internal cost of time [53–55].

The analysis of ground-reaction forces (GRFs) revealed increased vertical loading and lateral oscillations during co-manipulation. However, human-SLs instability was reduced by the mirror mode, where the second SL moved symmetrically to the one holding the object: the standard deviation of left–right GRF decreased by about 17% (≈ 30 N) relative to nominal control during co-manipulation, and total task duration also shortened. This supports coordinating multiple SLs at the coupled human–robot system level [2]: when the task SL induces inertial or balance perturbations, the other SL(s) can counterbalance to preserve stability while maintaining manipulation performance, thereby indirectly contributing to execution [6, 7]. More broadly, these results indicate that wearable SLs should be controlled as an internally coordinated ensemble rather than as independent effectors.

In conclusion, autonomous SLs can be safely and seamlessly integrated into human motor planning, enabling a single operator to perform complex, accuracy-constrained tasks that otherwise cannot be completed alone. By exploiting intrinsic haptics, legible motion timing, and mirrored multi-limb coordination, this work delineated behavioral and control principles—fluency, stability, and usability—that chart a path toward integrating SLs as practical tools across diverse applications from industrial operations to assistance for people with motor impairments [1, 2].

## 4 Methods

### 4.1 Participants and material

The protocol was approved by Imperial College London’s Science, Engineering and Technology Research Ethical Committee (SETREC approval number: 7111986). A total of 8 participants (1 female, 7 male), of age 27.3 ± 5.5 years old (mean std), height 176 ± 8.2 cm, weight 75.3± 9.4 kg, were recruited to carry out a tube assembly task with their arms and two wearable SLs. Each participant was provided an informed consent form, which they signed before performing the experiment.

*Dr. Hexapus* system used in our experiments consists of two 7-DoFs robotic arms (Gen3, Kinova, Boisbriand, Québec, Canada) mounted on a custom made backpack described in the Results section. These cobots comply with international safety standards for close interaction with humans [2]. Emergency stop buttons cutting the SLs power are available for the augmented participant, as described in the Results section. Each SL is equipped with a robotic gripper, resulting in an approximate weight of 8 kg per SL (see Fig. 1C for details), for a payload capacity of 4 kg per SL. The weight of the backpack and robot represented roughly 40% of the participants’ weight. The robots are controlled at 500 Hz, and joint torques and kinematics are recorded at 100 Hz.

This is part of the *MUltilimb Virtual Environment* (MUVE, see Supplementary Fig. S.1 and Fig. 2C) enables synchronous interaction with up to four of the SLs presented above. MUVE is equipped with a motion capture system (VICON, Oxford, UK) recording at 100 Hz. Here, the motion capture system was used to provide (i) the position and orientation of the PVC tube and the backpack in the task space, allowing to obtain the position and orientation of the SLs bases; and (ii) the human hands and head kinematics to build the safety constraints preventing collisions between SLs and the participant, and evaluate the effect of the different control modes on the participant’s movement. The platform also includes a treadmill equipped with force plates (GRAIL platform, Motek, Houten, Netherlands), yielding the *ground reaction force* (GRF) at 100 Hz, which was used to assess the effects of the different control modes on the participants’ balance.

### 4.2 Human-SLs collaborative task

Participants performed a dynamic task in collaboration with the SLs, which involved co-manipulating and assembling a long PVC tube (diameter 5 cm, length 3 m). They were instructed to: (i) grasp the tube with the right hand; (ii) raise it above the head; (iii) hand it over to the left hand; (iv) move it to the target location; and (v) assemble it by screw fastening while stabilizing with the left hand. Participants were briefed on the SLs’ general behavior–specifically, that approaching the tube with the right hand would initiate the task and that the left SL would be triggered when the left hand approached the tube during handover. They were not informed about the specific SL control strategies. From the SLs’ perspective, the task can be split into seven actions as follows (see Fig. 2A–C):

1. **Prepare right SL tube grasping:** The right SL moves from its home position to an intermediate goal, located at 10 cm from the tube axis with a final orientation constraint pointing the gripper towards the tube.
2. **Right grasping motion:** The right SL performs a linear motion to reach and grasp the tube.
3. **Tube co-manipulation:** The right SL co-manipulates the tube with the human’s right hand, to an intermediate goal above their head.
4. **Move left SL to prepare switching:** The left SL moves from their home position to an intermediate goal, located 10 cm from the tube axis, with a final orientation constraint pointing the gripper towards the tube.
5. **Left grasping motion:** The left SL performs a linear motion to reach and grasp the tube.
6. **Disengage right SL:** The right SL releases its grasp and performs a linear motion to disengage from the tube.
7. **Tube co-manipulation and assembly:** The left SL co-manipulates the tube with the human’s left hand to the final assembly target. The diameter of the target hole is approximately 5.5 cm. The tube is then stabilized while the human performs the assembly to the target using three screws.

Importantly, this task cannot be performed with only two human arms, as the assembly requires to simultaneously stabilize the tube and place screws to mount it on the target.

We tested two main control modes to implement the human-SLs collaboration. In the first, the *nominal control mode*, the seven actions were performed sequentially by the SLs, as described above. In the second, the *mirror mode*, the left SL was controlled to mirror the right SL movements during actions 1 and 2, and perform action 3 in parallel to the right SL. This control mode was tested because it allows the weight of the right SL to be counterbalanced, which could improve the user experience. For each of these two modes, we investigated two movement speeds, fast (close to the maximum velocity achievable with our cobots, identified during pretests where the safety constraints of the robots led to stopping before ending the movement) and slow (roughly 30% slower). This resulted in a total of four conditions: {*nominal slow* (NS), *nominal fast* (NF), *mirror slow* (MS), *mirror fast* (MF)}.

The above sequence of actions for the co-manipulation and assembly of the tube was repeated three consecutive times in each control mode and with each velocity. Between each control mode, participants were given five minutes rest period. After performing three trials with one of the SLs control modes and at the end of the experiment, participants were asked to rate the cognitive and physical ergonomics of the task on eight questions inspired by the NASA-TLX questionnaire [33]. For each item they had five choices ranging from “Totally disagree” to “Totally agree”. No difference between control modes was observed from the questionnaires filled at the end of each condition. The detailed questionnaire items filled at the end of the experiment are provided in the Supplementary Section 3.

### 4.3 Data analysis

Motion capture data included the participant’s head, hand and trunk (using markers on the backpack) and the tube. The latter were used to compute the tube velocities through numerical differentiation and were then processed using a low-pass filter (Butterworth, fourth order, 5 Hz cut-off frequency). The SLs EEF velocities were obtained by (i) extracting the pose of the robot base from motion capture data, (ii) extracting the SLs joints angles, (iii) applying forward kinematics, and (iv) filtering and numerically differentiating the resulting SLs EEF task space position profiles.

Average trajectories and velocity profiles for each subject, condition, and action were obtained by resampling all data at 1 kHz based on the average duration of the action across trials. Across population average profiles were obtained using the same method. The smoothness of the velocity profiles was assessed using the dimensionless jerk, computed using the accompanying codes from [32]. For all reported parameters, average values were computed for each subject, condition, and action across trials before plots.

To assess possible causal relationships between the human and SL movements, we used Granger causality tests [56]. This analysis was performed using the *grangercausalitytests* function from the *statsmodels*.*tsa*.*stattools* package, between the average SL EEF and tube velocity profiles obtained in action 3 & 7 for each condition. The significance level of the Granger tests was set at *p* = 0.05.

Given the size of our dataset, and the rejection of the normality hypothesis after performing Shapiro-Wilk tests [57], we used non-parametric statistical analyses to assess the effects of (i) the action performed by the human and SLs, and (ii) the tested control modes, on their respective behaviors. Main effects of action and control mode were assessed using Friedman tests, with a significant level set at *p* = 0.05.

When a significant main effect was detected, Wilcoxon signed-rank tests with a Benjamini-Hochberg false discovery rate correction were applied as post-hocs to assess differences between conditions. The significance level of these tests was set at *p* = 0.05, and we report Cohen’s *D* as a measure of the effect size.

### 4.4 SLs motion control

The control of the SLs follows a hierarchical implementation as described in Fig. 6. The top layer triggers each action based on the human and SLs states and the control mode, and extracts safety constraints at the level of the scene. Based on these states and constraints, the middle layer computes each SL’s action goal (a 6-dimensional pose) and plans an optimal trajectory to reach it, before the low-level executes the movement.

**Figure 6:**
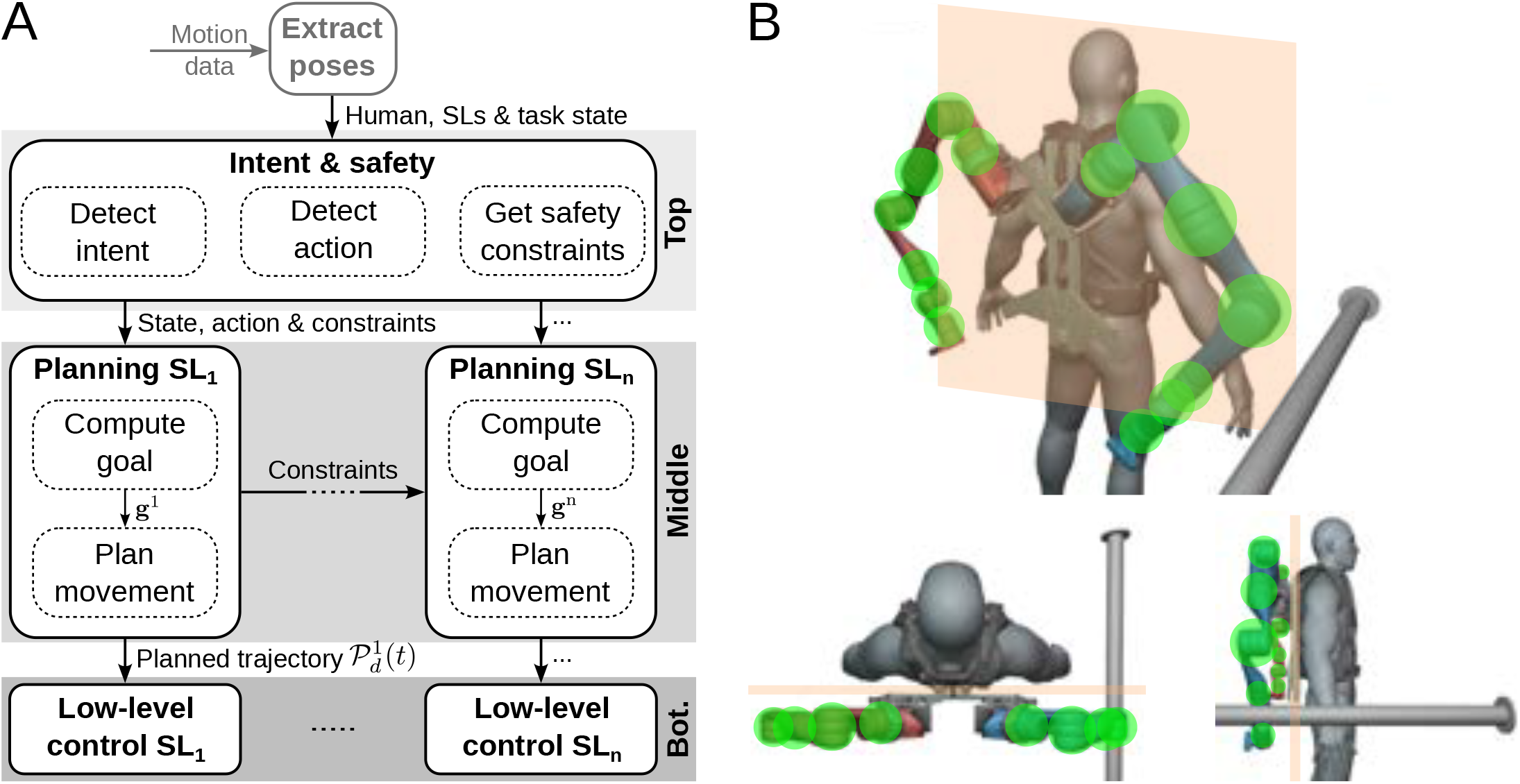
Hierarchical control framework for safe autonomous behaviors in humans-SLs collaboration. **A**. The framework allows to safely control n SLs by (i) detecting human intent, the task phase and generating safety constraints, (ii) using optimal motion planning under constraints to generate SLs trajectories, and (iii) implementing a low-level control of the SLs guaranteeing the execution of their motor plan. **B**. Illustration of the implemented safety constraints. The orange shaded rectangles represent the vertical virtual wall limiting the SLs operation zone. The green disks represent the spheres defined at the SLs joints.

#### Triggering of actions

Our control modes require the SLs to detect which action to perform. The task is described as a sequence of actions represented as a graph with a set of state transitions constraints *C*, similar to the approach used in [58, 59]. The graph representations of the action triggering mechanisms are provided in Fig. 2B for the two control modes, where each state label corresponds to the performed action, except for the initial and final states. Here we detail the implementation of the state transitions constraints, two of which trigger action sequences based on human behavior, while the others depend only on the SL states.

The first constraint 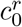 is verified when the human right hand comes close to the tube, which is when the distance between the hand and the tube axis (estimated using the motion capture system) falling below 10 cm. When verified, this triggers action 1, the initial movement of the SL.

In the nominal mode, constraint 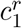 is verified when the right SL reaches the intermediate task goal 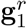, corresponding to the tube approach pose. Constraint 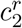 behaves similarly and is verified when the right SL has grasped the tube, i.e. when 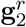 is reached. Constraint 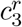 is verified when (i) the right SL has finished co-manipulating the tube, i.e. 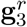 is reached; (ii) the tube position is above the participant’s head, meaning that the closest point of the tube to the head is higher than the head markers; (iii) both of the participant’s hands are close to the tube, i.e. the minimal distance between each hand and the tube axis is below 10 cm. Finally, *c*_4_, *c*_5_, *c*_6_, and *c*_7_ are successively verified when the relevant SL finishes its planned movement, corresponding to reaching 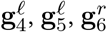 and 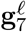.

In mirror mode, 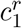 and 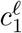 are verified when both SLs have completed the first planned movement, reaching 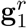 and 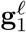. Similarly, 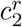 and 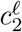 are verified when the SLs reach their respective goals 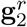 and 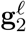, and the tube is grasped by the right SL. Then, 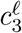 is verified when 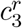 is satisfied and the left SL has completed its approach motion and reached 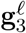. Finally, *c*_4_ does not exist in mirror mode, and the remaining constraints are identical to the nominal mode.

##### Defining goals

As transitions between task actions often rely on the SLs reaching their goal in task space, extracting these goals during human-SL collaboration is required for a successful task completion. In our setup, all goals were computed from motion capture data during task execution, adapting to each participant’s anthropometrics and posture and ensuring safe, reachable targets.

A major requirement was to ensure that the linear tube grasping motions of actions 2 & 5 were successful. First, at the end of actions 1 & 4, the goal poses 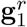 and 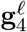 (or 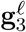 in mirror mode) constrained the SL gripper to lie in the (**z, x**) plane and to be perpendicular to the tube axis, respectively (see Fig. 2C for the axes definition). Second, the EEFs goal positions were placed at 10 cm from the tube axis in polar coordinates, with respective approach angles of −45^◦^ for 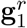 and 90^◦^ for 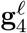 (or 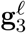 in the mirror mode). Finally, the corresponding linear movements were performed 30 cm and 45 cm behind the user’s back for actions 2 & 5, respectively, to avoid any collision with the human user.

Next, we ensured that the transfer of the tube from the right to the left SL was performed above the user’s head to avoid any collision between the tube and the head. Accordingly, the goal pose 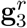 was set 30 cm above the head markers, with the SL gripper vertical and contained in the (**z, x**) plane of the world frame. The **y** components of 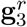 and 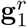 were set equal so that the co-manipulation movement remained mostly planar in the (**x, z**) plane.

Finally, the last goal pose ensured correct positioning of the tube in front of the target for assembly. The orientation of 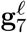 was constrained so that the tube axis was collinear with the normal to the target plane and passing through the center of the target hole. Rotation of the tube around its own axis remained unconstrained to allow the user to align the screw holes.

#### Planning of SLs movements

The method used to plan SL movements depended on the type of action performed. First, the small movements required to reach and grasp the tube in actions {2,5,6} were generated as straight movements in task space with a smooth bell-shaped velocity profile minimizing the jerk [60, 61] and lasting 2 s. In the absence of mechanical coupling with the human, during a *tube approach* (actions 1 & 4), we planned optimal SL movements without explicitly accounting for human biomechanical constraints. Conversely, during *co-manipulation* (actions 3 & 7), it was critical to ensure that the trajectories followed by the SLs considered the human arms biomechanics. Otherwise, the planned SL motion, which constrains the human through the tube, could result in movements impossible to perform or uncomfortable for the user. Below, we describe (i) the general control strategy and (ii) the different cost terms and differences between the tube approach and co-manipulation cases.

##### General motion planning strategy

In both the *tube approach* and *co-manipulation* cases, the task consisted of moving either the right or left SL EEF from an initial goal pose **g**_*i*−1_ ∈ ℝ^6^ to a final goal pose **g**_*i*_ ∈ ℝ^6^, while avoiding obstacles and remaining within the SL joints limits and velocity capabilities. The state of an SL at instant *t* is denoted by 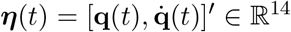, where 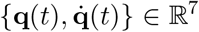 represent the SL joints positions and velocities, and ^′^ denotes the transpose operator.

To obtain feasible, collision-free and smooth SL movements in *tube approach* or *co-manipulation*, we minimized the following common cost function using a *Gaussian process motion-planner* (GPMP2) [62,63] and the Levenberg-Marquardt algorithm [64],

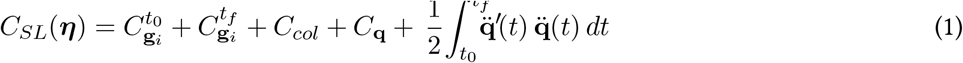

where 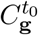 and 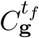 enforce the initial and final poses, *C*_col_ penalizes collisions along the generated path, *C*_q_ penalizes violations of joint limits and velocity constraints, and the last term penalizes the magnitude of joint acceleration 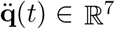 to ensure smooth movements. In GPM2, this cost corresponds to the *prior cost* and is implemented as a penalty on deviations from a constant velocity model. Below, we detail the implementation of each cost term.

##### Enforcing goal poses

Deviations from the desired initial pose are penalized in joint space:

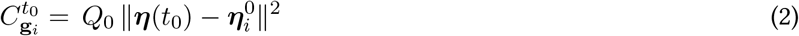

where 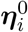 is the initial desired state corresponding to **g**_*i*−1_, and *Q*_0_ = 0.5 10^6^ the associated weight. Deviations from the desired final pose are penalized both in joint and task spaces as follows,

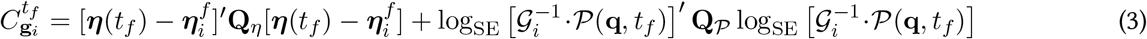

where 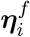 is the desired final state associated with **g**_*i*_ and **Q**_***η***_ ∈ ℝ^14×14^ penalizes only deviations from the desired joint velocities with a weight of 10^6^, for the joint space part. At time *t*_*f*_, deviations of the SL EEF pose *P*(*t*_*f*_) from the goal pose **g**_*i*_, with associated matrix G_*i*_, are penalized in task space using the *log* in SE, i.e. “boxminus” operator, defined in ℝ^6^ in [65] (p.52, Eq. 10.7). The weight matrix **Q**_*p*_ = 10^4^ diag(1, 10^−2^, 1, 1, 10^−2^, 1) scales the cost of the final pose deviations. Note that deviations along and around the **y**-axis are reduced as the problem is largely invariant to translation along the tube axis (mostly aligned with **y**) and rotation around it.

##### Obstacles avoidance

Throughout the task execution, it was critical to ensure that the considered SL did not collide with any obstacles, namely the human, the tube (when it was not grasped) or the other SL.

To efficiently represent obstacles, the task space (2 m × 3.5 m × 2.5 m in the **x, y, z** frame) was discretized into 6 cm × 6 cm × 6 cm voxels labeled as “occupied” or “free” depending on whether any obstacle intersected them.

Both SLs were covered by *K* spheres **s**_*k*_ with centers spaced 9 cm apart and radius *r*_*k*_ = 5 cm (so that adjacent spheres overlapped) to represent the arm segments. Spheres with larger radius *r*_*k*_ = 7.5 cm were used at the joints and EEF (see Fig. 6B for an illustration). The task-space positions of the sphere centers were computed using the robot’s forward kinematics. A collision was detected when the distance between a sphere center and an obstacle was smaller than the sphere radius.

A *signed distance field* (SDF) was used to represent obstacle distances. It was stored as a vector containing the distances between the center of each sphere of the considered SL and each occupied voxel **d**_*col*_ ∈ ℝ^*D*^, with *D* = *K O* where *O* denotes the number of occupied voxels. The SDF was computed at each planning timestep for every sphere of the controlled SL.

The cost associated with obstacle avoidance was

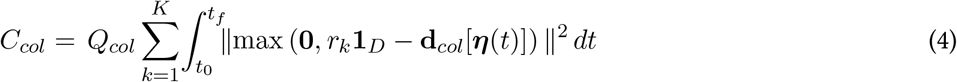

where *t*_0_ and *t*_*f*_ denote the initial and final times of the movement, and **1**_*D*_ ∈ ℝ^*D*^ is a vector of ones. The element-wise max function imposes a cost only in the case of penetration. The collision weight was set to *Q*_*col*_ = 2 10^6^, ensuring that colliding trajectories were always suboptimal.

To avoid collisions with the human, we constructed a vertical wall of occupied voxels in the task space parallel to the human coronal plane (see Fig. 6B for an illustration), computed using reflective markers placed on the backpack. This wall served as a task-space boundary that none of the spheres representing the SLs could cross. Finally, during *tube approach* movements, the tube was represented by its axis and radius.

##### Accounting for joints limits

The joints limits in position and velocity were constrained by minimizing:

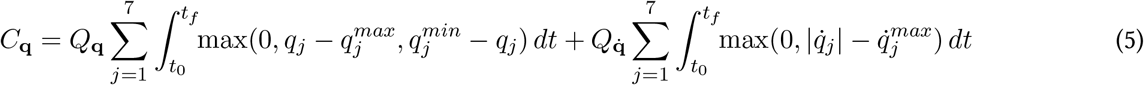

where *j* ∈ [[1, 7]] denotes an SL joint, 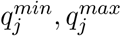 are the angle limits of joint *j* and 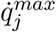 its maximal velocity. The associated weights were set as 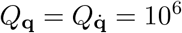.

##### Tube approach case

In absence of mechanical coupling between the human hands and the SLs, the cost function to minimize was defined as in Eq. 1. The initial guess trajectory was a constant-velocity linear interpolation between the initial ***η***_0_ and final ***η***_*i*_ states. The final state ***η***_*i*_ corresponded to the goal pose *G*_*i*_, and was computed using an iterative damped least-square inverse kinematics solver with random initializations for robustness [66, 67].

##### Co-manipulation case

When the manipulated object induced mechanical coupling, the SL movements also needed to remain compatible with human biomechanics to ensure feasible and comfortable physical interaction. For instance, in action 3 the SLs should perform outward-curved movements (see average trajectories in Fig. 2D). In contrast, inward-curved movements would impose large internal/external shoulder rotations of the shoulder during arm elevation together with full elbow flexion, resulting in excessive constraints on the user’s joints. For the two actions (3 & 7) involving tube co-manipulation, an initial desired pose trajectory with outward curvature and compatible with the obstacles was generated using cubic splines, which were adapted before acton execution due to the online computation of movement goals. The cost function of Eq. 1 was then extended with a tracking term enforcing the desired trajectory:

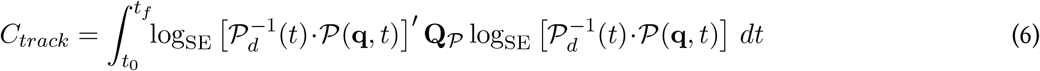

where *P*_*d*_(*t*) denotes the desired pose trajectory and **Q**_P_ is defined as in Eq. 3.

#### 4.4.1 Low-level control

Accurate execution of the desired SL trajectories was achieved using a *proportional-integral-derivative* (PID) correction scheme in joint space, made possible by computing joint-space trajectories via inverse kinematics. The gains of the PID corrector were

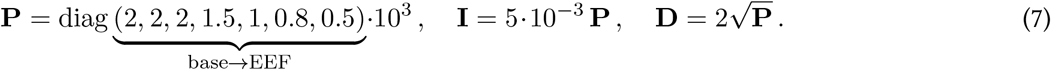

Higher values were used for joints closer to the base to (i) compensate for the greater effort required due to the weight and inertia of the SLs, and (ii) allow the wrists joints to remain slightly compliant, enabling small deviations from the desired pose during interaction with the human. The gains were selected so that the SLs could track the trajectories computed using the optimal planner while avoiding any control instability.

## Supporting information

Explanatory video

## Data availability statement

All data supporting the findings of this study are available within the paper, and can be provided upon request to the corresponding author.

## Acknowledgments

This study was funded in part by the UK EPSRC (MUVE EP/W036495/1) and the European Commission (FETOPEN H2020 899626 NIMA, ICT 871767 ReHyb, ERC 810346 NaturalBionics). Dorian Verdel was funded in part by the Agence Nationale de la Recherche, France 2030 program, reference PEPR O2R – PI3 ASSISTMOV (22-EXOD-0004).

## Contribution statement

- Conceptualization – DV, BX, AD, LM, PS, EB
- Methodology – DV, BX, HCC, AD, LM, EB
- Software – DV, BX, HCC, AD
- Validation – DV, AD
- Formal analysis – DV, BX, LM, AD
- Investigation – BX, HCC
- Resources – DF, EB
- Data Curation – DV, AD
- Visualization – DV, AD, EB
- Supervision – DV, HCC, EB
- Project administration – DF, EB
- Funding acquisition – DF, EB
- Writing - Original Draft – DV, AD, DF, EB
- Writing - Review & Editing – all authors

## References

[1] J. Eden, M. Bräcklein, J. Ibáñez, D. Y. Barsakcioglu, G. Di Pino, D. Farina, E. Burdet, and C. Mehring, “Principles of human movement augmentation and the challenges in making it a reality,” Nature Communications, vol. 13, no. 1, p. 1345, 2022.

[2] D. Verdel, J. Eden, H. Cervantes Culebro, M. Pinardi, G. Di Pino, P. Souères, C. Mehring, and E. Burdet, “A predictive coding framework for safe and versatile control of supernumerary robotic limbs,” The International Journal of Robotics Research, pp. 1–21, 2026, In press.

[3] D. Prattichizzo, M. Pozzi, T. L. Baldi, M. Malvezzi, I. Hussain, S. Rossi, and G. Salvietti, “Human augmentation by wearable supernumerary robotic limbs: review and perspectives,” Progress in Biomedical Engineering, vol. 3, p. 042005, sep 2021.

[4] B. Yang, J. Huang, X. Chen, C. Xiong, and Y. Hasegawa, “Supernumerary robotic limbs: a review and future outlook,” IEEE Transactions on Medical Robotics and Bionics, vol. 3, pp. 623–639, Aug. 2021.

[5] A. Dragan and S. Srinivasa, “Integrating human observer inferences into robot motion planning,” Autonomous Robots, vol. 37, pp. 351–368, aug 2014.

[6] C. Moon and J. Kim, “Strategies for moment compensation in supernumerary robotic limbs manipulation tasks,” in 2024 33rd IEEE International Conference on Robot and Human Interactive Communication (ROMAN), pp. 491–496, IEEE, Aug. 2024.

[7] X. Qiu, D. Verdel, H. Cervantes-Culebro, A. Devillard, and E. Burdet, “Augmenting human balance with generic supernumerary robotic limbs,” arχiv, 2026.

[8] B. L. Bonilla and H. H. Asada, “A robot on the shoulder: coordinated human-wearable robot control using coloured petri nets and partial least squares predictions,” in IEEE International Conference on Robotics and Automation (ICRA), pp. 119–125, may 2014.

[9] J. Zhang, H. Zeng, J. Liu, D. Chen, and A. Song, “Adaptive human movement compensation control of supernumerary robotic limb for overhead support task with non-zero-sum differential game theory,” IEEE Transactions on Automation Science and Engineering, vol. 22, pp. 14158–14169, 2025.

[10] J. Zhang, J. Zhang, H. Zeng, D. Chen, H. Li, and A. Song, “Development and control of supernumerary robotic limbs for overhead tube manipulation task,” IEEE Robotics and Automation Letters, vol. 11, pp. 2634–2641, Mar. 2026.

[11] F. Y. Wu and H. H. Asada, ““hold-and-manipulate” with a single hand being assisted by wearable extra fingers,” in 2015 IEEE International Conference on Robotics and Automation (ICRA), pp. 6205–6212, IEEE, May 2015.

[12] I. Hussain, G. Salvietti, G. Spagnoletti, and D. Prattichizzo, “The soft-sixthfinger: a wearable emg controlled robotic extra-finger for grasp compensation in chronic stroke patients,” IEEE Robotics and Automation Letters, vol. 1, no. 2, pp. 1000–1006, 2016.

[13] A. Shafti, S. Haar, R. Mio, P. Guilleminot, and A. A. Faisal, “Playing the piano with a robotic third thumb: assessing constraints of human augmentation,” Scientific Reports, vol. 11, Nov. 2021.

[14] K. Umezawa, Y. Suzuki, G. Ganesh, and Y. Miyawaki, “Bodily ownership of an independent supernumerary limb: an exploratory study,” Scientific Reports, vol. 12, Feb. 2022.

[15] D. Clode, L. Dowdall, E. da Silva, K. Selén, D. Cowie, G. Dominijanni, and T. R. Makin, “Evaluating initial usability of a hand augmentation device across a large and diverse sample,” Science Robotics, vol. 9, May 2024.

[16] F. Parietti and H. H. Asada, “Supernumerary robotic limbs for aircraft fuselage assembly: body stabilization and guidance by bracing,” in IEEE International Conference on Robotics and Automation (ICRA), pp. 1176–1183, may 2014.

[17] F. Parietti and H. Asada, “Supernumerary robotic limbs for human body support,” IEEE Transactions on Robotics, vol. 32, pp. 301–311, apr 2016.

[18] Stelarc, “Writing one word with three hands simultaneously.” url: stelarc.org/, 1982.

[19] N. Yamamura, D. Uriu, M. Muramatsu, Y. Kamiyama, Z. Kashino, S. Sakamoto, N. Tanaka, T. Tanigawa, A. Onishi, S. Yoshida, S. Yamanaka, and M. Inami, “Social digital cyborgs: the collaborative design process of JIZAI ARMS,” in Proceedings of the 2023 CHI Conference on Human Factors in Computing Systems, ACM, apr 2023.

[20] E. Abdi, E. Burdet, M. Bouri, S. Himidan, and H. Bleuler, “In a demanding task, three-handed manipulation is preferred to two-handed manipulation,” Scientific Reports, vol. 6, Feb. 2016.

[21] U. Koike, G. Enriquez, T. Miwa, H. E. Yap, M. Kabasawa, and S. Hashimoto, “Development of an intraoral interface for human-ability extension robots,” Journal of Robotics and Mechatronics, vol. 28, pp. 819–829, Dec. 2016.

[22] Y. Huang, E. Burdet, L. Cao, P. T. Phan, A. M. H. Tiong, and S. J. Phee, “A subject-specific four-degree-of-freedom foot interface to control a surgical robot,” IEEE/ASME Transactions on Mechatronics, vol. 25, pp. 951–963, apr 2020.

[23] G. Salvietti, I. Hussain, D. Cioncoloni, S. Taddei, S. Rossi, and D. Prattichizzo, “Compensating hand function in chronic stroke patients through the robotic sixth finger,” IEEE Transactions on Neural Systems and Rehabilitation Engineering, vol. 25, no. 2, pp. 142–150, 2016.

[24] S. Gurgone, D. Borzelli, P. de Pasquale, D. J. Berger, T. L. Baldi, N. D’Aurizio, D. Prattichizzo, and A. d’Avella, “Simultaneous control of natural and extra degrees of freedom by isometric force and electromyographic activity in the muscle-to-force null space,” Journal of Neural Engineering, vol. 19, p. 016004, jan 2022.

[25] G. Dominijanni, D. L. Pinheiro, L. Pollina, B. Orset, M. Gini, E. Anselmino, C. Pierella, J. Olivier, S. Shokur, and S. Micera, “Human motor augmentation with an extra robotic arm without functional interference,” Science Robotics, vol. 8, Dec. 2023.

[26] T. Lisini Baldi, N. D’Aurizio, C. Gaudeni, S. Gurgone, D. Borzelli, A. d’Avella, and D. Prattichizzo, “Exploiting body redundancy to control supernumerary robotic limbs in human augmentation,” The International Journal of Robotics Research, Aug. 2024.

[27] D. Leal Pinheiro, J. Faber, S. Micera, and S. Shokur, “Neuromuscular learning and control of auricular muscles for human–machine interfaces,” The International Journal of Robotics Research, Nov. 2025.

[28] B. Llorens-Bonilla, F. Parietti, and H. H. Asada, “Demonstration-based control of supernumerary robotic limbs,” in IEEE/RSJ International Conference on Intelligent Robots and Systems (IROS), oct 2012.

[29] E. Burdet, “MUltilimb Virtual Environment (MUVE) platform.” url: imperial.ac.uk/human-robotics/dr-octopus-/, 2023.

[30] https://www.motekmedical.com/solution/grail/.

[31] T. Yoshikawa, “Manipulability of robotic mechanisms,” The International Journal of Robotics Research, vol. 4, pp. 3–9, June 1985.

[32] S. Balasubramanian, A. Melendez-Calderon, A. Roby-Brami, and E. Burdet, “On the analysis of movement smoothness,” Journal of NeuroEngineering and Rehabilitation, vol. 12, pp. 1–11, dec 2015.

[33] S. G. Hart and L. E. Staveland, “Development of NASA-TLX (task load index): results of empirical and theoretical research,” in Advances in Psychology, pp. 139–183, Elsevier. 1988.

[34] J. Luo, X. Zhou, Y. Zhu, Y. Li, C. Zhang, K. Wang, Z. Mai, and C. Zeng, “A human–robot collaboration control framework for supernumerary robotic limbs,” Journal of Field Robotics, vol. 43, pp. 34–48, July 2025.

[35] P. Kieliba, D. Clode, R. O. Maimon-Mor, and T. R. Makin, “Robotic hand augmentation drives changes in neural body representation,” Science Robotics, vol. 6, may 2021.

[36] C. Mehring, M. Akselrod, L. Bashford, M. Mace, H. Choi, M. Blüher, A.-S. Buschhoff, T. Pistohl, R. Salomon, A. Cheah, O. Blanke, A. Serino, and E. Burdet, “Augmented manipulation ability in humans with six-fingered hands,” Nature Communications, vol. 10, jun 2019.

[37] M. Hao, J. Zhang, K. Chen, H. Asada, and C. Fu, “Supernumerary robotic limbs to assist human walking with load carriage,” Journal of Mechanisms and Robotics, vol. 12, July 2020.

[38] Z. Tu, Y. Jiang, H. Yan, Y. Leng, and C. Fu, “Design, modeling, control, and evaluation of a wearable Centaur robot for load-carriage walking assistance,” The International Journal of Robotics Research, Feb. 2026.

[39] Z. Zhao, S. Cheng, Y. Ding, Z. Zhou, S. Zhang, D. Xu, and Y. Zhao, “A survey of optimization-based task and motion planning: from classical to learning approaches,” IEEE/ASME Transactions on Mechatronics, vol. 30, pp. 2799–2825, Aug. 2025.

[40] M. G. Mohanan and A. Salgoankar, “A survey of robotic motion planning in dynamic environments,” Robotics and Autonomous Systems, vol. 100, pp. 171–185, Feb. 2018.

[41] J. J. Kuffner and S. M. LaValle, “RRT-connect: an efficient approach to single-query path planning,” in IEEE International Conference on Robotics and Automation., vol. 2, pp. 995–1001, 2000.

[42] E. Todorov and W. Li, “A generalized iterative LQG method for locally-optimal feedback control of constrained nonlinear stochastic systems,” in Proceedings of the American Control Conference., pp. 300–306, 2005.

[43] Y. Wen and P. Pagilla, “Path-constrained and collision-free optimal trajectory planning for robot manipulators,” IEEE Transactions on Automation Science and Engineering, vol. 20, pp. 763–774, Apr. 2023.

[44] M. Khoramshahi, A. Poignant, G. Morel, and N. Jarrassé, “A practical control approach for safe collaborative supernumerary robotic arms,” in IEEE Conference on Advanced Robotics and its Social Impact (ARSO), pp. 147–152, 2023.

[45] O. Khatib, “Real-time obstacle avoidance for manipulators and mobile robots,” The International Journal of Robotics Research, vol. 5, pp. 90–98, Mar. 1986.

[46] M. Zucker, N. Ratliff, A. D. Dragan, M. Pivtoraiko, M. Klingensmith, C. M. Dellin, J. A. Bagnell, and S. S. Srinivasa, “CHOMP: covariant hamiltonian optimization for motion planning,” The International Journal of Robotics Research, vol. 32, pp. 1164–1193, Aug. 2013.

[47] M. Pinardi, M. R. Longo, D. Formica, M. Strbac, C. Mehring, E. Burdet, and G. Di Pino, “Impact of supplementary sensory feedback on the control and embodiment in human movement augmentation,” Communications Engineering, vol. 2, Sept. 2023.

[48] A. Takagi, G. Ganesh, T. Yoshioka, M. Kawato, and E. Burdet, “Physically interacting individuals estimate the partner’s goal to enhance their movements,” Nature Human Behaviour, vol. 1, pp. 1–6, Mar. 2017.

[49] L. E. Miller, L. Montroni, E. Koun, R. Salemme, V. Hayward, and A. Farnè, “Sensing with tools extends somatosensory processing beyond the body,” Nature, vol. 561, pp. 239–242, Sept. 2018.

[50] J. W. Guggenheim and H. H. Asada, “Inherent haptic feedback from supernumerary robotic limbs,” IEEE Transactions on Haptics, vol. 14, pp. 123–131, jan 2021.

[51] G. Dominijanni, S. Shokur, G. Salvietti, S. Buehler, E. Palmerini, S. Rossi, F. De Vignemont, A. d’Avella, T. R. Makin, D. Prattichizzo, and S. Micera, “The neural resource allocation problem when enhancing human bodies with extra robotic limbs,” Nature Machine Intelligence, vol. 3, pp. 850–860, Oct. 2021.

[52] M. Pinardi, A. Noccaro, L. Raiano, D. Formica, and G. Di Pino, “Comparing end-effector position and joint angle feedback for online robotic limb tracking,” PLOS ONE, vol. 18, p. e0286566, June 2023.

[53] R. Shadmehr, J. J. O. de Xivry, M. Xu-Wilson, and T.-Y. Shih, “Temporal discounting of reward and the cost of time in motor control,” Journal of Neuroscience, vol. 30, pp. 10507–10516, aug 2010.

[54] B. Berret and G. Baud-Bovy, “Evidence for a cost of time in the invigoration of isometric reaching movements,” Journal of Neurophysiology, vol. 127, pp. 689–701, feb 2022.

[55] D. Verdel, O. Bruneau, G. Sahm, N. Vignais, and B. Berret, “The value of time in the invigoration of human movements when interacting with a robotic exoskeleton,” Science Advances, vol. 9, sep 2023.

[56] C. W. J. Granger, “Investigating causal relations by econometric models and cross-spectral methods,” Econometrica, vol. 37, p. 424, Aug. 1969.

[57] S. S. Shapiro and M. B. Wilk, “An analysis of variance test for normality (complete samples),” Biometrika, vol. 52, p. 591, dec 1965.

[58] L. Johannsmeier and S. Haddadin, “A hierarchical human-robot interaction-planning framework for task allocation in collaborative industrial assembly processes,” IEEE Robotics and Automation Letters, vol. 2, pp. 41–48, Jan. 2017.

[59] K. Darvish, E. Simetti, F. Mastrogiovanni, and G. Casalino, “A hierarchical architecture for human–robot cooperation processes,” IEEE Transactions on Robotics, vol. 37, pp. 567–586, Apr. 2021.

[60] P. Morasso, “Spatial control of arm movements,” Experimental Brain Research, vol. 42, pp. 223–227, Apr. 1981.

[61] T. Flash and N. Hogan, “The coordination of arm movements: an experimentally confirmed mathematical model,” The Journal of Neuroscience, vol. 5, pp. 1688–1703, jul 1985.

[62] J. Dong, M. Mukadam, F. Dellaert, and B. Boots, “Motion planning as probabilistic inference using gaussian processes and factor graphs,” in Robotics: Science and Systems XII, 2016.

[63] M. Mukadam, J. Dong, X. Yan, F. Dellaert, and B. Boots, “Continuous-time gaussian process motion planning via probabilistic inference,” The International Journal of Robotics Research, vol. 37, pp. 1319–1340, Sept. 2018.

[64] H. P. Gavin, “The Levenberg-Marquardt algorithm for nonlinear least squares curve-fitting problems,” Department of Civil and Environmental Engineering Duke University August, vol. 3, no. 1-23, p. 4, 2019.

[65] J. L. Blanco-Claraco, “A tutorial on SE(3) transformation parameterizations and on-manifold optimization,” arχiv, 2021.

[66] J. Angeles, “On the numerical solution of the inverse kinematic problem,” The International Journal of Robotics Research, vol. 4, pp. 21–37, June 1985.

[67] S. R. Buss and J.-S. Kim, “Selectively damped least squares for inverse kinematics,” Journal of Graphics Tools, vol. 10, pp. 37–49, Jan. 2005.

