## Supplementary materials for "Out-of-sight partners: autonomous wearable supernumerary limbs for dynamic co-manipulation"

### 1 MUVE platform

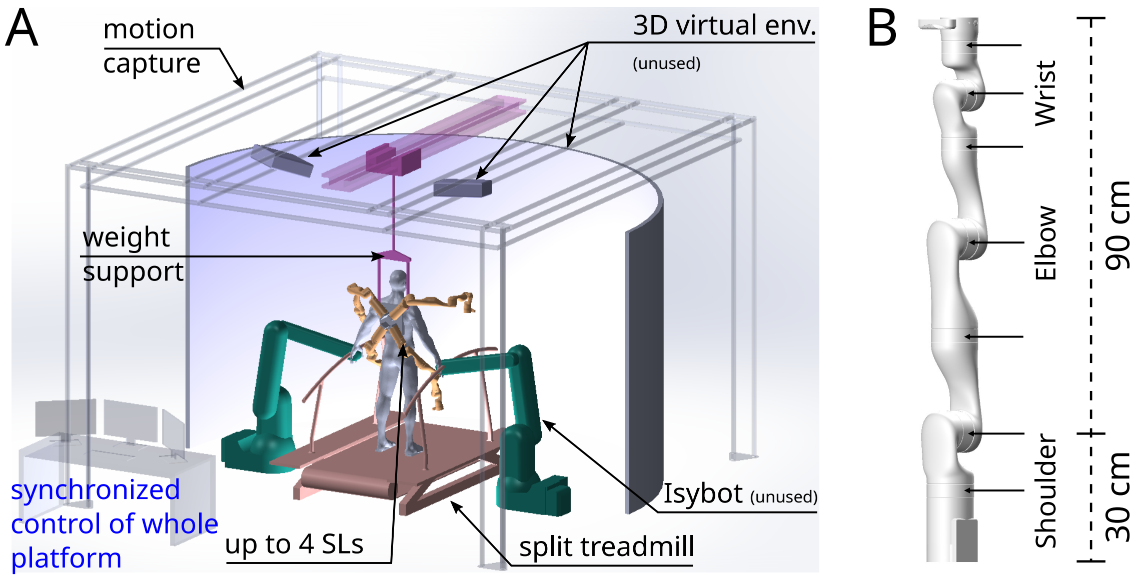


**Figure S.1: Experimental setup for augmentation research. A.** The MUlti-limb Virtual Environment (MUVE) is a full-body, multisensory (visual, auditory, haptic) platform enabling physical interaction with SLs in dynamic virtual environments. MUVE integrates: (i) 3D motion capture; (ii) a split treadmill with force plates; (iii) up to four lightweight, wearable robotic arms for movement augmentation (iv) immersive 3D VR (unused in the present work); and (v) two SYB3 (Isybot, Les Ulis, France) high-power robotic interfaces (unused in the present work). All components are synchronized and integrated within a single system using ROS. **B.** The selected SL with seven joints and dimensions allowing back-mounting, providing a workspace that overlaps the operator's natural reach for co-manipulation while extending posteriorly to access otherwise unreachable regions.

### 2 General SLs behavior

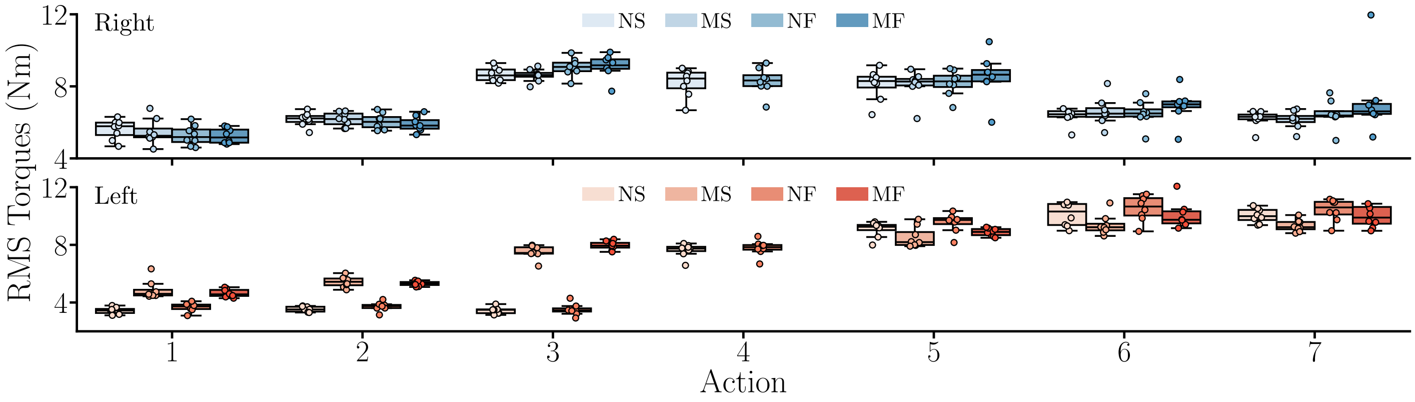


**Figure S.2: RMS of summed SLs' joints torques throughout task execution.** Data related to the right SL are represented in blue and to the left SL in red. Results are provided for the four tested conditions, i.e. nominal and mirror slow {NS,MS} and fast {NF,MF}.

Fig. 2E in the main text shows how the SLs increased their velocity between *slow* {NS,MS} and *fast* {NF,MF} conditions for actions 1, 3, 4 (in nominal mode) and 7. This was confirmed by a significant main effect of condition during actions 1 & 3 for the right SL (in both cases: Friedman's $W > 0.67$, $p \leq 10^{-3}$) and during actions 1–4 & 7 for the left SL ($W > 0.72$, $p < 0.001$ in all cases). In detail, slow conditions yielded lower right EEF velocities than fast conditions in actions 1 & 3 (in all cases: $p < 0.023$, Cohen's $D > 1.57$), and similarly lower left EEF velocities in actions 1–3 (mirror mode), 4 (nominal mode) & 7 ($p < 0.023$, $D > 1.53$ in all cases). As expected, in actions 1–3 the mirror produced higher left EEF velocities than the nominal mode ($p < 0.012$, $D > 4.91$ in all cases). In sum, participant movements of the participants and human-SL co-manipulation of the tube did not hinder the SLs' ability to modulate speed. No velocity difference were observed in actions 2, 5 & 6, as anticipated, because these approach/retraction phases followed the same linear motion plan.

Fig. S.2C shows that the RMS of torque across the seven joints of each SL depended primarily on the posture, the required movement, and on the interaction with the human. In the mirror mode, we observed a main effect of action on torques for both the right SL ($W = 0.82$, $p < 0.001$) and the left SL ($W = 0.96$, $p < 0.001$). For the right SL, co-manipulation/handover actions 3 & 5 elicited higher torques than all other actions ($p < 0.02$, $D > 2.65$ in all cases). No significant differences emerged among actions 2, 6 & 7, where the right SL was static or executing approach/retraction motions in the upper workspace. Action 1–free SL movements in the lower workspace–yielded significantly lower torques than all other actions ($p < 0.04$, $D > 1.56$ in all cases). For the left SL, torques increased across successive actions up to actions 6 & 7 ($p < 0.02$, $D > 1.42$ for all preceding pairwise comparisons), which did not differ significantly. Overall, tube co-manipulation phases (actions 3–5 for the right SL and 6 & 7 for the left SL) produced the highest torques, even in similar workspace regions without co-manipulation.

### 3 Questionnaire

Please tick the box that better describes your perception after interacting with the robotic system in the four blocks.

1. In all conditions, performing the task was physically tiring.
   - ☐ Totally agree
   - ☐ Somewhat agree
   - ☐ Neither agree nor disagree
   - ☐ Somewhat disagree
   - ☐ Totally disagree
2. In all conditions, performing the task was mentally tiring.
   - ☐ Totally agree

☐ Somewhat agree
☐ Neither agree nor disagree
☐ Somewhat disagree
☐ Totally disagree

3. In all conditions, I the task was simple to perform.

☐ Totally agree
☐ Somewhat agree
☐ Neither agree nor disagree
☐ Somewhat disagree
☐ Totally disagree

4. In all conditions, I could comfortably perform the task.

☐ Totally agree
☐ Somewhat agree
☐ Neither agree nor disagree
☐ Somewhat disagree
☐ Totally disagree

5. In all conditions, wearing the robots was comfortable.

☐ Totally agree
☐ Somewhat agree
☐ Neither agree nor disagree
☐ Somewhat disagree
☐ Totally disagree

6. In all conditions, I felt high pressure on my shoulders.

☐ Totally agree
☐ Somewhat agree
☐ Neither agree nor disagree
☐ Somewhat disagree
☐ Totally disagree

7. In all conditions, I was able to understand the robots behaviour.

☐ Totally agree
☐ Somewhat agree
☐ Neither agree nor disagree
☐ Somewhat disagree
☐ Totally disagree

8. In all conditions, I felt safe while interacting with the robots.

☐ Totally agree
☐ Somewhat agree
☐ Neither agree nor disagree
☐ Somewhat disagree
☐ Totally disagree
